# NLRP1 Shapes Immune and Inflammatory Signatures in Human Melanoma but Not in Mouse Models

**DOI:** 10.1101/2025.11.24.690170

**Authors:** Vitor Pedro Targhetta, Raquel de Souza Vieira, Hedden Ranfley, Cristiane Naffah de Souza Breda, Niels Olsen Saraiva Câmara

## Abstract

Inflammasomes are multiprotein complexes that activate pro-caspase-1, leading to the maturation of the pro-inflammatory cytokines IL-1β and IL-18. *Nlrp1*, the first receptor identified with inflammasome-forming capacity, is highly expressed in both the skin and immune cells. Despite its prominent role in these tissues, the function of *Nlrp1* in melanoma remains poorly characterized. In this study, we investigated the impact of *Nlrp1* on melanoma patient survival and found that its expression is associated with improved prognosis and with a co-expression network enriched for pro-inflammatory genes. However, in murine models, neither *Nlrp1* expression nor activation significantly affected melanoma development or progression. Similarly, pharmacological activation of *Nlrp1* using Val-Boro-Pro (VbP) did not alter tumor growth or the local inflammatory profile in mice but directly influenced CD25^+^ cell generation and glucose uptake in *in vitro* models. Finally, we demonstrated that a melanoma risk score can be constructed based on genes specific to inflammasome and pyroptosis pathways. Collectively, our findings reveal species-specific differences in NLRP1 function between humans and mice and support the potential of inflammasome-related pathways as prognostic biomarkers and therapeutic targets in human cancers.

## Introduction

Inflammasomes are cytosolic multiprotein complexes that function as innate immune sensors, initiating inflammatory responses upon the recognition of pathogen- or danger-associated molecular patterns (1,2). Their activation results in the cleavage of pro-caspase-1 into its active form, caspase-1, which subsequently promotes the maturation of interleukins IL-1β and IL-18 and induces pyroptosis—a lytic form of programmed cell death mediated by gasdermin-D cleavage (3).

Among the known inflammasome sensors, NLRP1 (NLR family pyrin domain containing 1), first described in 2002, was the earliest identified receptor capable of initiating inflammasome assembly (4). Its activation involves a complex sequence of proteolytic events and can be triggered by viral proteases, the anthrax lethal factor (LF), and the DPP8/9 inhibitor Val-Boro-Pro (VbP) (5–7).

NLRP1 is expressed across a wide range of tissues; however, RNA-seq data indicate that the skin exhibits the highest transcript levels. In keratinocytes, NLRP1 is the predominant inflammasome receptor, with expression levels far exceeding those of NLRP3, AIM2, or NLRC4, and it mediates cytokine production in response to UVB radiation (8,9). Gain-of-function variants in NLRP1 have been associated with several inflammatory skin disorders, including multiple self-healing palmoplantar carcinoma (MSPC), chronic lichenoid keratosis, vitiligo, and psoriasis (8, 10, 11). These associations highlight the pivotal role of NLRP1 in cutaneous immunity and inflammatory pathology.

Despite its established role in skin inflammation, the contribution of NLRP1 to cutaneous tumorigenesis remains less clearly defined. In vitro studies suggest that NLRP1 may exert pro-tumorigenic effects in melanoma, being associated with increased tumor cell viability, enhanced IL-1β secretion, NF-κB activation, and inhibition of caspase-mediated apoptosis. Consistently, gain-of-function NLRP1 variants in humans have been linked to an elevated susceptibility to skin tumors, likely through epithelial hyperplasia driven by the paracrine signaling of IL-1β, IL-1α, and IL-18 (8,12,13).

Given its high expression in the skin and its capacity to modulate both inflammatory responses and epithelial cell biology, NLRP1 may play a significant role in melanoma development, acting through both tumor-intrinsic and microenvironmental mechanisms. In this study, we sought to investigate the role of NLRP1 in melanoma progression and patient survival using complementary approaches in murine models and human datasets.

## Methodology

### Animals

All animal experiments were conducted in accordance with Federal Law No. 6,638 of 1979, which regulates the use of animals in scientific experimentation. The study was approved by the Ethics Committee on Animal Research of the University of Sa o Paulo – USP, Institute of Biomedical Sciences (CEUA No. 2906200220). Wild-type C57BL/6 (H-2^b^) and NLRP1 knockout (NLRP1 KO, all paralogous genes) male and female mice, aged 8–12 weeks, were obtained from the Animal Facility of the Department of Immunology, University of Sa o Paulo. Mice were maintained under controlled conditions with a 12-hour light/dark cycle, constant ambient temperature of 22 °C, and free access to autoclaved food and water.

### CD4^+^ T Cell Isolation and Polarization

Spleens and lymph nodes (periaortic, submandibular, subaxillary, and inguinal) were collected from mice, mechanically dissociated, and filtered through 70 µm cell strainers. CD4^+^ T lymphocytes were isolated using the EasySep™ Mouse CD4^+^ T Cell Isolation Kit (STEMCELL™ Technologies), following the manufacturer’s instructions. Isolated lymphocytes were seeded in 96-well plates pre-coated with anti-CD3 and cultured in RPMI medium supplemented with glutamine, pyruvate, essential amino acids, and penicillin/streptomycin. For Treg differentiation, cells were cultured in the presence of TGF-β (5 ng/mL), anti-IFN-γ (10 µg/mL), anti-IL-4 (10 µg/mL), anti-IL-12 (10 µg/mL), and anti-CD28 (1 µg/mL). The differentiation protocol was performed for 6 days at 37 °C.

### Staining and Flow Cytometry

Cells obtained from WT or NLRP1 KO mice were plated in V-bottom 96-well plates. After centrifugation, cells were resuspended in FACS buffer (1× PBS + 2% FBS) and incubated with surface antibodies for 30 minutes at 4 °C. Subsequently, cells were fixed and permeabilized using the Foxp3/Transcription Factor Staining Buffer Kit (TONBO Biosciences™) for intracellular staining of transcription factors and/or cytokines.

Flow cytometry acquisition was performed using a FACSCanto II cytometer (BD Biosciences) with FACSDiva software, and data were analyzed using FlowJo software (Tree Star, San Carlos, CA, USA).

### Subcutaneous Melanoma Model and VbP Treatment

The melanoma cell line used for animal injections was B16-F10 (ATCC® CRL-6475). These cells were cultured in complete high-glucose DMEM medium, as specified by ATCC. For the melanoma model, a suspension containing 2 × 10⁵ viable B16-F10 cells in serum-free high-glucose DMEM was subcutaneously inoculated into the dorsal region of WT and NLRP1 KO mice.

Tumor growth was monitored every two days after tumor onset (day 10) by measuring tumor dimensions with a caliper. Tumor volume (mm³) was calculated using the formula: Tumor Volume = (d² × D) × 0.5, where *d* represents the smaller diameter and *D* the larger diameter of the tumor mass.

For VbP treatment, mice received daily oral gavage of 20 µg VbP per animal for a period of one week.

### Acquisition and Analysis of TCGA Data

Raw data from the TCGA-SKCM cohort (FPKM, count tables, and clinical data) were obtained using the R package TCGAbiolinks and primarily used for the construction of the proposed gene signature. Genes with FPKM = 0 in more than half of the patients were excluded from the analysis.

The TIMER database was used to perform the univariate Cox regression analysis and to generate plots relating inflammasome- and pyroptosis-associated genes with the magnitude of B lymphocyte infiltration and patient survival (14). To correlate NLRP1 expression with different lymphocyte subtypes, data from the TISIDB repository were used (15). Differences in NLRP1 expression between primary and metastatic tumors were obtained from the UALCAN database (16).

The LinkedOmics platform was employed to construct the co-expression network of genes associated with NLRP1 and to perform pathway enrichment analysis of the resulting gene set (17).

### Construction of the Risk Score

Briefly, normalized expression data (FPKM) were obtained from the TCGA-SKCM cohort, along with patient survival information. The variable “days_to_death” was considered as the survival time; in cases where this variable was unavailable, “days_to_last_follow_up” was used instead. For the selection of genes evaluated, those belonging to the REACTOME INFLAMMASOMES (https://www.gsea-msigdb.org/gsea/msigdb/cards/REACTOME_INFLAMMASOMES) and REACTOME PYROPTOSIS (https://www.gsea-msigdb.org/gsea/msigdb/cards/REACTOME_PYROPTOSIS.html) pathways were included. After obtaining the expression matrices and identifying the key genes associated with inflammasomes and pyroptosis, we employed the R package glmnet to perform LASSO-Cox regression using the selected genes. This penalized regression approach refines the final model by performing variable shrinkage within the Cox proportional hazards framework, thereby reducing the number of predictors required to assess patient survival.

Survival curves were generated using the R packages survival and survminer, while ROC curves and their corresponding area under the curve (AUC) values were computed using the survivalROC package.

### Acquisition of Immunotherapy-Related Datasets and FASTQ Files

When necessary, raw FASTQ files were downloaded directly from the Sequence Read Archive (SRA – NCBI) using the SRA Explorer tool (https://sra-explorer.info/). The quality of the sequencing reads was assessed using FastQC, and when required, read trimming and quality adjustments were performed with the Trimmomatic software to ensure compliance with the desired quality standards.

Sequence alignment was performed using the HISAT2 software with default parameters and the pre-built Homo sapiens GRCh38 index (http://daehwankimlab.github.io/hisat2/download/) (18). The resulting .sam files were converted to .bam format using the Samtools utility (19). The aligned reads, together with the reference annotation (*H. sapiens* GRCh38 v104), were used for transcript assembly with STRINGTIE, and all assembled transcripts were subsequently merged into a pooled annotation to generate an updated reference containing novel transcripts derived from individual samples (20). Using this new reference annotation, the aligned reads were then used for transcript quantification. A count table was generated using the prepDE.py script provided with STRINGTIE. The gffcompare tool was employed to compare the reference annotation with the newly assembled transcript annotation.

### Statistical Analysis

In animal experiments, data are presented as mean ± standard deviation. All datasets were assumed to follow a Gaussian distribution with homogeneity of variance. Comparisons between two groups were performed using an unpaired, two-tailed Student’s *t*-test. For comparisons involving more than two groups, one-way ANOVA was applied, followed by Fisher’s LSD post hoc test when appropriate for multiple comparisons. Cox proportional hazards regression analysis was used for survival analyses, unless otherwise specified. A significance level (*α*) of 0.05 was considered for all statistical tests.

## Results

### High expression of NLRP1 is related to longer survival and immune infiltration in patients with melanoma

Data related to melanoma from the TCGA portal (TCGA-SKCM) were analyzed to initially assess whether NLRP1 expression influences melanoma progression. To begin our analyses, we examined the correlation between patient survival and NLRP1 expression in the TCGA-SKCM cohort using the TIMER database. Patients were stratified according to the median NLRP1 expression level and subsequently evaluated for overall survival. The results reveal a significantly longer survival among patients exhibiting higher NLRP1 expression (Figures 1A and B).

**Figure 1.**
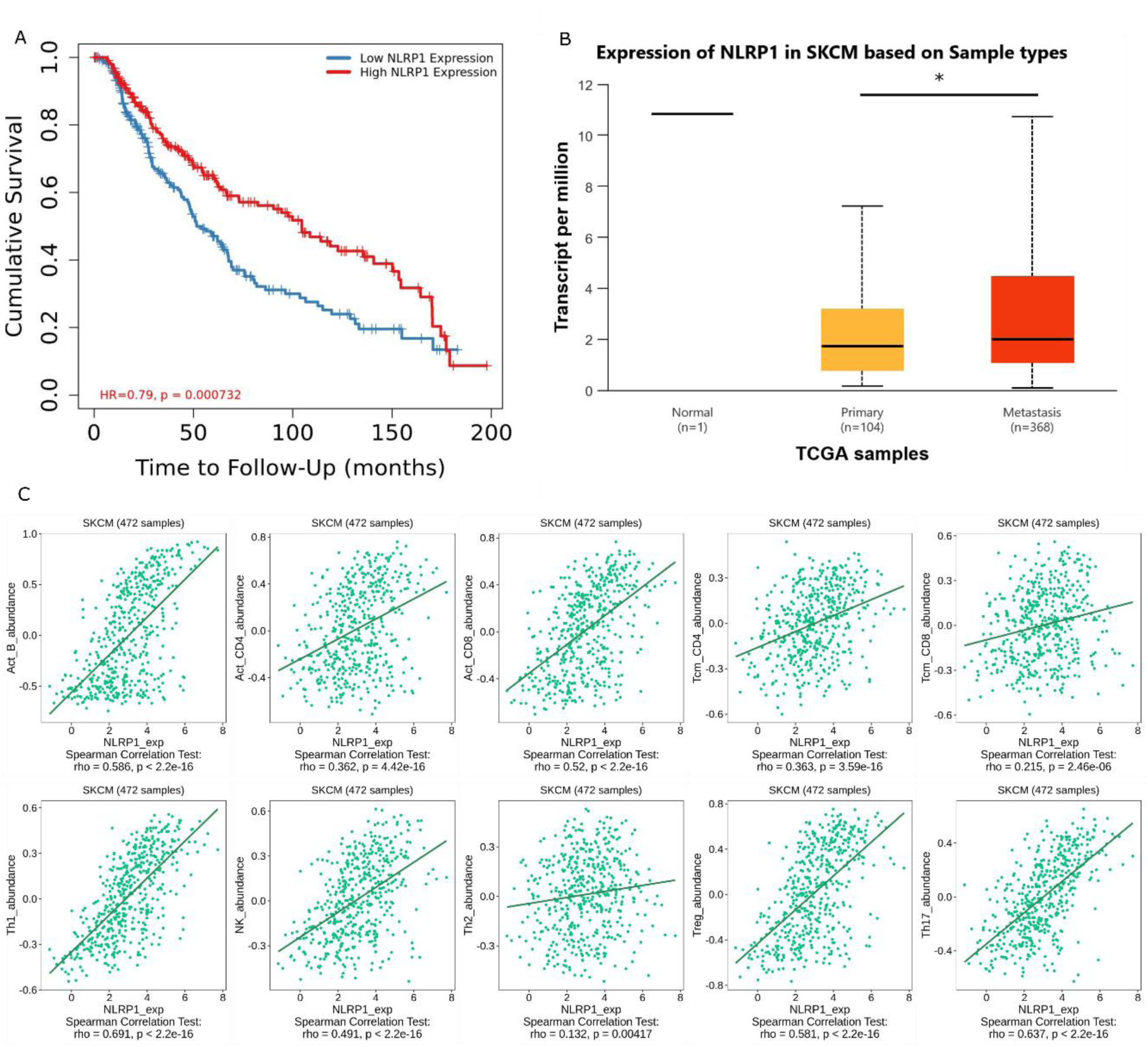
NLRP1 correlates with improved survival and immune infiltration in melanoma. **A.** Using the TIMER database, patients were stratified according to median of NLRP1 expression levels, and survival analyses were performed using a univariate Cox model, with overall survival plotted in months. **B.** TCGA-SKCM patient data were further stratified by tumor subtype and NLRP1 expression levels (in TPMs) using the UALCAN repository. **C.** Correlation between NLRP1 expression and immune cell subtypes in the TCGA-SKCM cohort is shown. The TISIDB database was used to perform Spearman correlation analyses with major immune subtypes derived from deconvolution of the TCGA-SKCM transcriptomic data.

Furthermore, analysis using the TISIDB repository demonstrated a positive correlation between NLRP1 expression and the principal adaptive immune cell types within the tumor microenvironment (Figure 1C). Collectively, these findings indicate that NLRP1 expression is positively associated with both improved overall survival and increased immune cell infiltration, suggesting that its role in the tumor microenvironment is, at least in part, mediated through modulation of immune infiltrates.

### NLRP1 co-expressed genes are associated with a pro-inflammatory phenotype in human melanoma

To understand the potential mechanisms through which NLRP1 may contribute to the regulation of tumor evolution in melanoma, we analyzed the gene coexpression network associated with NLRP1 and the main biological processes involved. For this purpose, we utilized the coexpression functional module of the LinkedOmics database to assess RNA-seq data from the TCGA-SKCM cohort, identifying 4,105 genes whose expression was significantly correlated with that of NLRP1 (Figure 2A).

**Figure 2.**
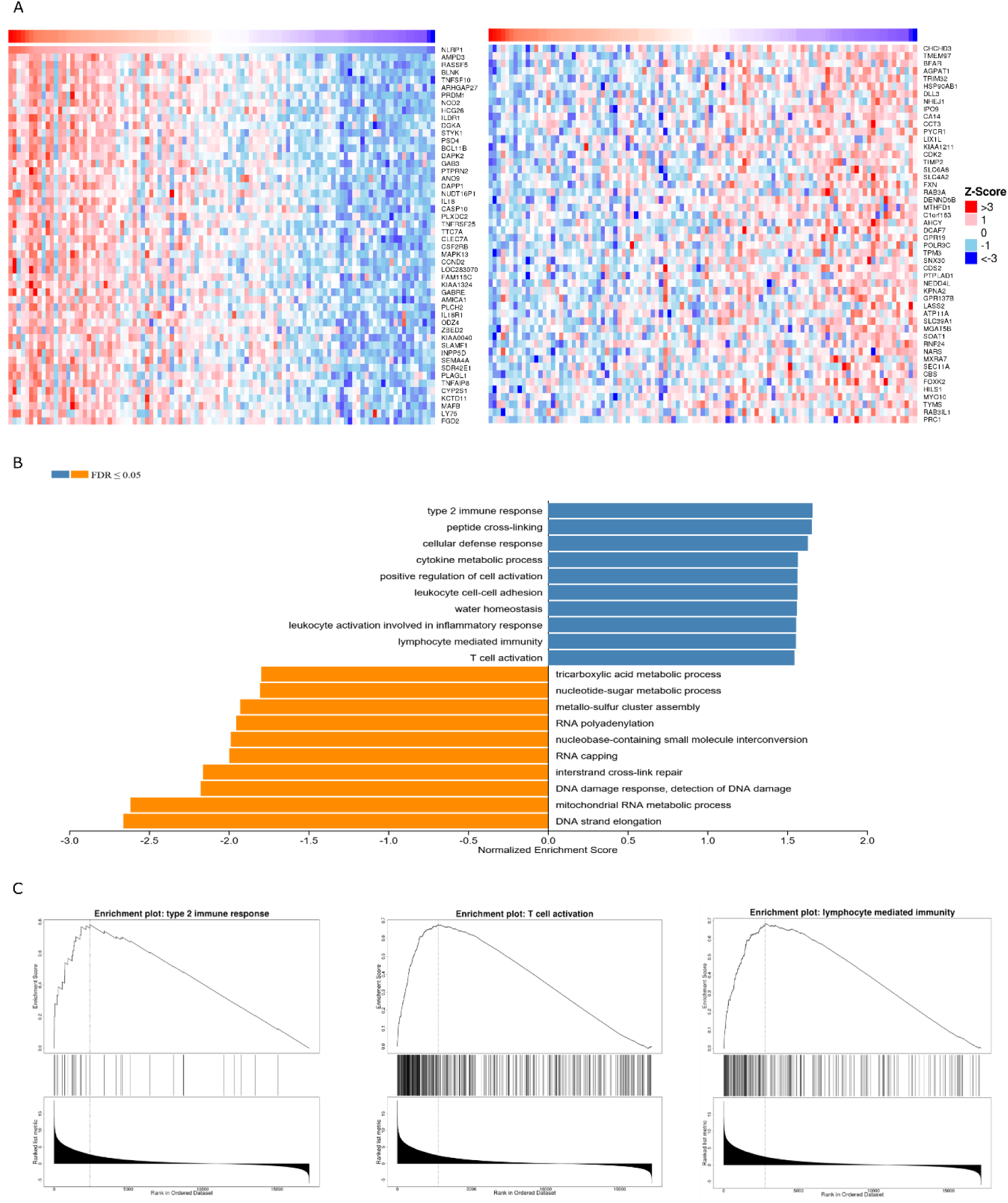
The NLRP1 co-expression network in melanoma points to pro-inflammatory phenotypes. **A.** Heatmap showing the top 50 genes positively and negatively correlated with NLRP1 expression, respectively. Heatmaps were obtained from the LinkedOmics repository, with gene expression values normalized to Z-scores (mean = 0, standard deviation = 1). The 50 most significantly correlated genes are shown—positively correlated on the left and negatively correlated on the right. **B.** Representative GSEA plots of genes positively and negatively associated with NLRP1. Analyses were performed using the LinkedOmics platform and plotted according to pathway enrichment scores and FDR values. **C.** Representative GSEA plots of three biological processes positively associated with NLRP1 expression.

To further characterize these results, a gene set enrichment analysis (GSEA) was performed using genes that were positively (ρ > 0.3) and negatively (ρ < –0.3) correlated with NLRP1 expression, as determined by the LinkedOmics database. The analysis revealed that positively correlated genes were predominantly enriched in pathways related to adaptive immune maintenance and T cell activation—most notably, the “Type 2 immune response,” “interleukin-4 production” and “adaptive immune response” pathways (Figures 2 B and C). Conversely, genes negatively correlated with NLRP1 were primarily enriched in pathways associated with transcription, translation, and RNA/DNA metabolism (Figure 2B).

### The presence and activation of NLRP1 does not influence the development of melanoma in mice

High NLRP1 expression has previously been associated with promoting tumor growth in melanoma by enhancing cell survival, as well as contributing to tumorigenesis in other cancer types, such as breast cancer. However, little is known about its role within the tumor microenvironment or whether these effects are directly dependent on NLRP1 activation.

To assess the importance of NLRP1 in tumor control—and given its apparent relevance to human survival—we compared the tumor growth of the B16-F10 murine melanoma cell line in C57BL/6 and NLRP1 knockout (KO) mice. The results revealed no significant differences in melanoma growth between the groups evaluated (Figures 3A and B). Tumor size did not differ between the two mouse strains, suggesting that the basal antitumor response to subcutaneous melanoma development is likely independent of NLRP1.

**Figure 3.**
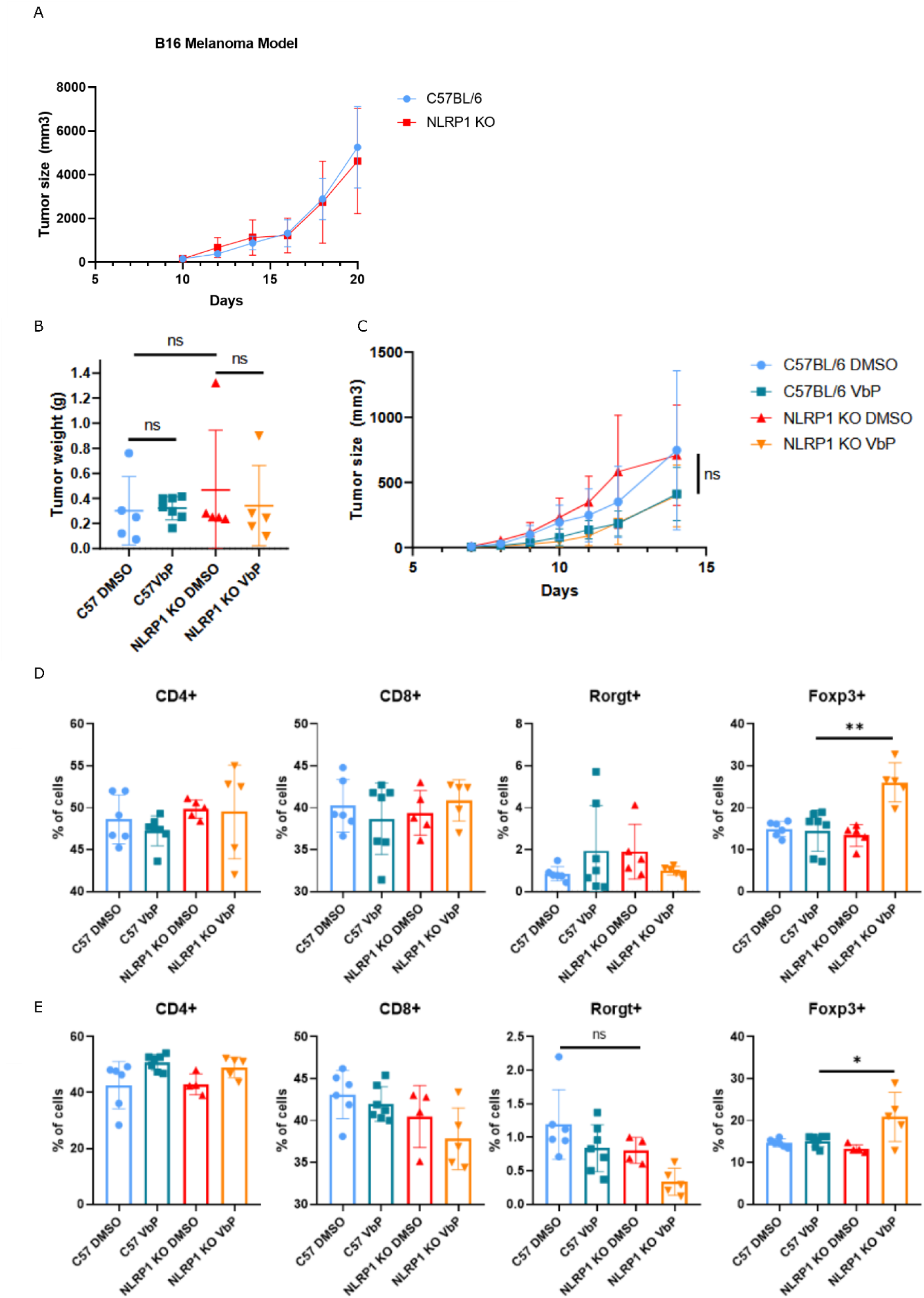
Melanoma model in WT and NLRP1 KO mice following VbP treatment. **A.** Tumor growth curve of the subcutaneous melanoma model in C57BL/6 and NLRP1 KO mice. Mice were inoculated intradermally with 2×10⁵ B16-F10 cells and monitored for 20 days to assess tumor growth during this period. **B.** B16 tumor weights from mice treated with VbP or vehicle control (DMSO) by oral gavage after 15 days. **C.** Mean tumor growth curves comparing the experimental groups from B. **D-E.** Phenotyping of CD8^+^ and CD4^+^ T lymphocytes and their Th17 and Treg subsets in tumor-draining lymph nodes (D) or contralateral lymphonodes (E) from mice treated or not with VbP.

Next, we sought to determine whether pharmacological activation of NLRP1 by VbP could confer therapeutic benefits by delaying melanoma progression. VbP has previously been shown to exert potent antitumor effects in murine bladder carcinoma models and to enhance the efficacy of anti–PD-1 treatment in pancreatic cancer models. To test this, we subcutaneously injected B16-F10 cells into C57BL/6 and NLRP1 KO mice and after tumor establishment, animals were treated orally with VbP for seven consecutive days (Figure 3C). Our results indicate that VbP treatment did not significantly delay melanoma growth, and no differences were observed in the frequencies of total CD4^+^ or CD8^+^ T cells in VbP-treated animals compared with their respective controls, both in draining (Figure 3D) and contralateral (Figure 3E) lymph nodes.

### VbP impacts CD25+ cell generation independently of NLRP1

Given the intricate relationship between NLRP1 and the immune system in humans, we also compared the global immune phenotypes of C57BL/6 and NLRP1 KO mice (Supplementary Figure 1). No significant differences were observed in the proportions of the major immune cell subsets. Interestingly, however, the presence of VbP was found to reduce the differentiation of regulatory T cells (CD4^+^Foxp3^+^) in vivo and to decrease the percentage of CD4^+^CD25^+^ cells among splenocytes, in a manner independent of NLRP1 (Figure 4A). Additionally, VbP treatment reduced glucose uptake in splenocytes, as indicated by a lower percentage of 2NBDG-positive cells (Figure 4B). These findings suggest a potential new mechanism of action for VbP, whereby both the inhibition of glucose uptake and the reduction in CD25^+^ cell generation may contribute to its antitumor effects. Despite altering the abundance of glucose in the intracellular environment, no differences were observed in the percentage of mitochondria among the experimental groups (Figure 4C).

**Figure 4.**
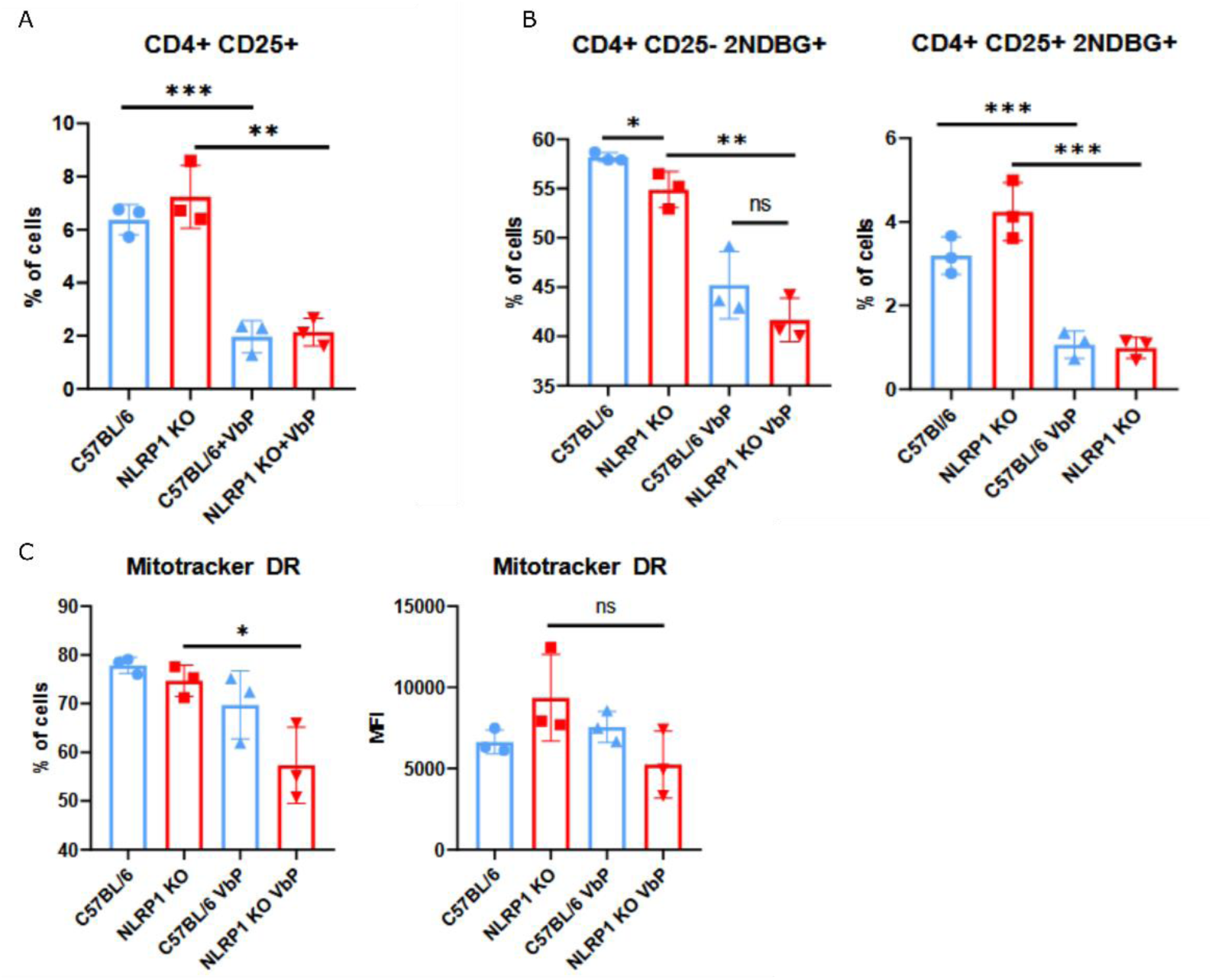
VbP impacts CD25^+^ cell generation and glucose uptake. **A.** CD25 expression in CD4^+^ T lymphocytes following VbP treatment. Total splenocytes from C57BL/6 and NLRP1 KO mice were treated with 20 µM VbP for 24 hours and subsequently analyzed for the percentage of CD25^+^ cells within the CD4^+^ T cell population. **B.** Splenocytes from were treated with 20 µM VbP or DMSO for 24 hours, then incubated with 2-NBDG (25 µM) and stained for surface markers to assess total CD4^+^ T cells and Tregs (CD25^+^). **C.** Mitochondrial polarization assay using MitoTracker Deep Red (50 nM) after treatment with 20 µM VbP or DMSO for 24 hours. Legend: p < 0.05; **p** < 0.01; \***p** < 0.001.

### NLRP1 is not directly related to a better response to immunotherapy in patients

To assess if the described effect of enhancing anti-PD1 immunotherapy of VbP is NLRP1 dependent, we decided to verify whether the high expression of NLRP1 could also have beneficial effects in anti PD-1 immunotherapy. To begin our analysis, we collected studies in the GEO-NCBI database (Gene Expression Omnibus) that contained deposited RNAseq datasets referring to melanoma patients treated with anti-PD-1 antibody, together with data referring to the RECIST criterion of response to immunotherapy. Two datasets were found, GSE78220 and GSE91061, in which tumor biopsies were sequenced prior to treatment, which presented a good number of samples and were within the initial criteria evaluated (Supplementary Table 1). We compared NLRP1 expression in responders (Complete Response and Partial Response) and non-responders (Stable Disease and Progressive Disease) where no initial difference was observed (Supplementary Figure 2A). Additionally, in the evaluated data, NLRP1 was not able to predict patient survival (Supplementary Figure 2B). These results suggests that NLRP1 expression is not directly relevant to immunotherapy success.

### Expression of genes correlated with inflammasome and pyroptosis can build a risk score for melanoma patients

We found that not only NLRP1, but others inflamassome and pyroptosis related genes have a strong tendency to be positively correlated with longer survival of patients with melanoma in the TCGA-SKCM cohort (Supplementary Figure 3). To quantify these events, we evaluated the proportional risk of each gene based on a univariate Cox-type linear regression (Univariate Cox hazard ratio – Cox HR) and the results of the significant genes in patient survival are presented in the figure below (Figure 5A).

**Figure 5.**
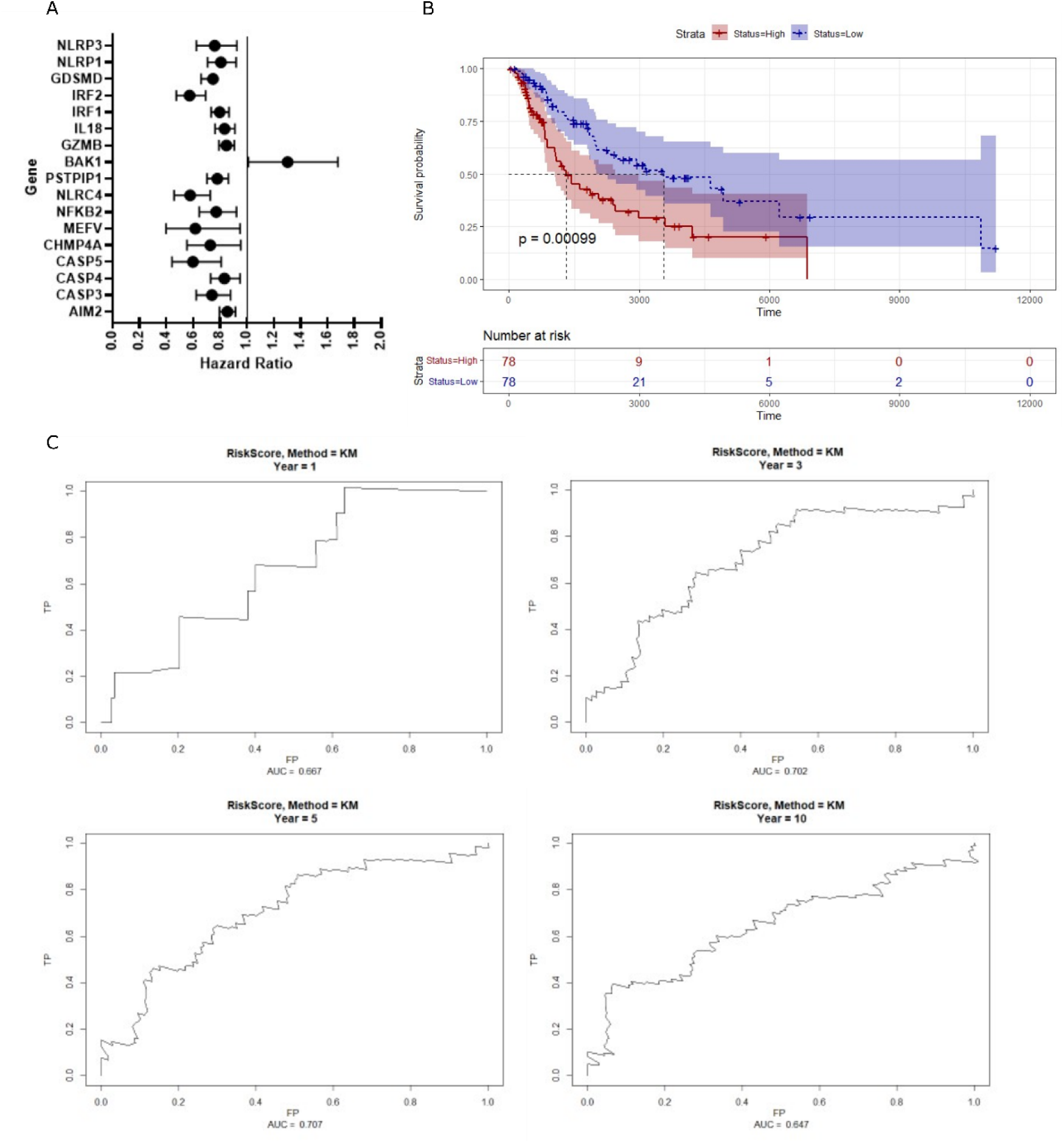
Construction of a risk score based on inflammasome- and pyroptosis-related genes. **A.** Seventeen genes associated with patient survival were identified with a significant hazard ratio (HR ≠ 1) in the TCGA-SKCM cohort. **B.** Survival curve based on the median value of the proposed risk score. After calculating the risk score for each patient in the test subcohort, patients were divided according to the median risk value, and the corresponding risk score (Cox-HR) was used to generate a Kaplan–Meier survival curve showing survival distribution over time. **C.** ROC curves and corresponding areas under the curve (AUC) representing 1-, 3-, 5-, and 10-year survival in the test cohort based on the constructed risk score model.

Of the inflammasome and pyroptosis genes tested, 17 correlated significantly with survival. Of these genes, in 16 the high expression correlated directly with better survival (HR < 1) and in only 1 gene, BAK1, patients who expressed it less had better survival (HR > 1). The observation that a significant portion of genes related to inflammasomes and pyroptosis can be a good predictive clinical marker of tumor evolution in melanomas led us to try to build a risk score model based on this type of genetic signature. To accomplish this, we separated the TCGA-SKCM cohort into two distinct sub-cohorts: the training data (315 patients, corresponding to 2/3 of the total) and the test data (156 patients, the remaining 1/3). We trained our model in the training data using the linear regression model of the LASSO-Cox type from the clinical data and normalized expression in FPKM of the previously found significant genes for survival to build a risk score equation of the type: score = ∑ coefficient(x)*gene(x).

After selecting the minimum lambda for adjusting the regression (Supplementary Figure 4), we obtained the relative coefficients of each gene and apllied it to obtain the equation, resulting in: Risk score = (MEFV * 0.75) – (AIM2 * 0.0146) + (BAK1 * 0.0122) – (CASP4 * 0.00065) – (CASP5) * 0.328) – (CHMP4A * 0.1197) – (GSDMD * 0.01) – (GZMB * 0.00569) – (IRF1 * 0.03) – (IRF2 * 0.0286) + (NFKB2 * 0.0168) – (NLRC4 * 0.2058) + (NLRP3 * 0.1048) – (PSTPIP1 * 0.04).

To validate the previously obtained risk score, we measured it for each of the patients in the sub-cohort of the test model (156 patients) who were not part of its construction. After obtaining each respective score, we divided the patients according to the median and separated them into ’High’ (50% with higher scores) and ’Low’ (50% with lower scores). Interestingly, as a result, we obtained a significantly lower survival in patients in the ’High risk’ group, which demonstrates that the score was functional in predicting patients with a poor prognosis (Figure 5B). Additionally, the ROC curves related to patient survival within the previously proposed risk model, for 1, 3, 5 and 10 years, demonstrate that the risk score was functional in predicting patient survival (Figure 5C).

## Discussion

Only a few studies in the literature have attempted to associate NLRP1 with tumorigenesis, and among these, the findings reveal substantial heterogeneity regarding its role, which may be either pro- or anti-tumoral (13,21).

Our in-silico analyses demonstrated that NLRP1 expression correlates with an increased infiltration of lymphocytes within the tumor and with the co-expression of genes regulating T cell responses. Furthermore, NLRP1 was associated with improved overall survival in patients with melanoma. These findings contrast with previous in vitro studies and analyses of patients harboring NLRP1 gain-of-function variants, in which NLRP1 was primarily described as a pro-tumoral factor in melanoma (8,13,22,23). The activation of NLRP1 can trigger pyroptosis in human cell types such as keratinocytes, monocytes, and macrophages, which constitute a significant portion of the skin. Therefore, one might expect these cells to also play a relevant role in shaping the melanoma tumor microenvironment. However, our results revealed no significant differences in tumor growth between C57BL/6 and NLRP1 KO mice inoculated with the murine melanoma cell line B16. It is important to emphasize that these results primarily reflect effects within the tumor stroma, since only the host mice—but not the B16 cell line—lacked the NLRP1 gene. These findings also differ from observations regarding other inflammasomes, such as NLRP3. In NLRP3 KO mice, a significant reduction in pulmonary metastatic foci was reported in an intravenous B16 injection model (24). In this context, it is important to note that NLRP1 exhibits marked divergence between humans and mice. In the murine genome, three paralogous genes have been identified—Nlrp1a, Nlrp1b, and Nlrp1c, the latter being a pseudogene. Each of these genes displays substantial variability in both presence and nucleotide sequence among commonly used mouse strains (25). In contrast, humans possess a single NLRP1 gene. Nonetheless, although there are distinct activators for both proteins, the inhibition of DPP8/9 by Vbp was shown to be a universal activator of NLRP1 in humans and mice (5).

Our data also indicates that treatment with VbP did not reduce melanoma progression and did not alter lymphocyte infiltration, either in the draining lymph node or in the contralateral lymph node. This outcome contrasts with reports in which VbP has shown antitumor activity, either as a single agent or in combination with other therapies. The discrepancy between our results and previous findings likely reflects differences in tumor models. VbP was shown to be effective in murine models of acute myeloid leukemia (26), bladder tumors, rhabdomyosarcoma (27), and pancreatic cancer (28) —the latter being relatively less immunogenic, whereas the others are considered highly immunogenic models (29,30). Also, the antitumor effects of VbP were reported to be immune-dependent (27,28), suggesting that the low immunogenicity of the B16 melanoma model, combined with the high intratumoral cell death observed, may contribute to the lack of therapeutic response in our experiments.

The VbP treatment also reduced the frequency of CD4^+^ Foxp3^+^ Tregs. A concomitant reduction in CD25^+^ lymphocytes was also detected in splenocytes following VbP exposure. Since these effects occurred independently of NLRP1, we hypothesize that the direct enzymatic targets of VbP, namely DPPIV, DPP8, and DPP9, may mediate the observed outcomes. Among these, DPPIV is the most extensively characterized member of the dipeptidyl peptidase (DPP) family. It is a membrane-associated enzyme, ubiquitously expressed across most cell types, and plays a co-stimulatory role in T cell activation, particularly via the TCR signaling pathway, enhancing migration and proliferation of T cells (31). Despite its high expression in effector T cells (Th1 and Th17), DPPIV is considered a negative marker for Tregs, with virtually absent expression in this subset (32,33). Given its absence in Tregs and its localization at the plasma membrane, the most likely intracellular targets of VbP responsible for the observed reduction in Foxp3 expression are DPP8 and DPP9, two cytoplasmic enzymes with high sequence and structural similarity. Although their precise physiological functions remain largely undefined, both are expressed in lymphoid organs, including the spleen, thymus, and lymph nodes, supporting a potential role in immune regulation (32). The best-characterized biochemical function of DPP8 and DPP9 is the cleavage of N-terminal Xaa–Pro dipeptides (34). Notably, the Foxp3 protein contains three consecutive Xaa–Pro motifs within its N-terminal region, theoretically making it a potential substrate for these enzymes.

Finally, recent studies have proposed several inflammasome-related gene signatures associated with survival and tumor progression in cancers such as ovarian, cervical, and hepatocellular carcinoma, among others (35–39). These models accurately predicted patient survival within their respective cohorts, underscoring the relevance of pyroptosis and the secretion of IL-1β and IL-18, downstream products of inflammasome activation, in tumor development across diverse contexts. Our prognostic model was generated from an initial set of approximately 50 candidate genes derived from the REACTOME pyroptosis and inflammasome pathways. Genes with significant hazard ratios (n = 17) were selected, and 14 genes were ultimately retained in the final LASSO-Cox regression, providing a refined signature for predicting melanoma outcomes. Like those previously reported, our risk score also proved effective in predicting patient survival, indicating that the expression of genes related to pyroptosis and the inflammasome may serve as a basis for predictive analyses and, consequently, improve patient prognosis.

## Suplemmentary Files

**Sup. Table 1.** Characteristics of the datasets analyzed in relation to anti–PD-1 resistance.

| Characteristic | Hugo et al. (GSE78220) | Riaz et al. (GSE91061) |
| --- | --- | --- |
| Number of samples | 28 | 33 |
| Number of responders | 15 | 10 |
| Number of non-responders | 13 | 23 |
| Biopsy timing | Pre-treatment | Pre-treatment |
| Previous treatment | Variable (MAPKi) | Variable (ipilimumab) |
| Mean age | 60 years | Not available |
| Sex | Male/Female | Male/Female |
| Drug administered | Nivolumab/Pembrolizumab | Nivolumab |
| RNA selection | Poly(A) tail | Poly(A) tail |
| Sequencing platform | Illumina HiSeq 2000 | Illumina Genome Analyzer |
| Sequencing strategy | Paired-end | Paired-end |
| Read length | 2 × 100 bp | 2 × 51 bp |
| Strand specificity | Unstranded | Unstranded |

**Sup. Figure 1.**
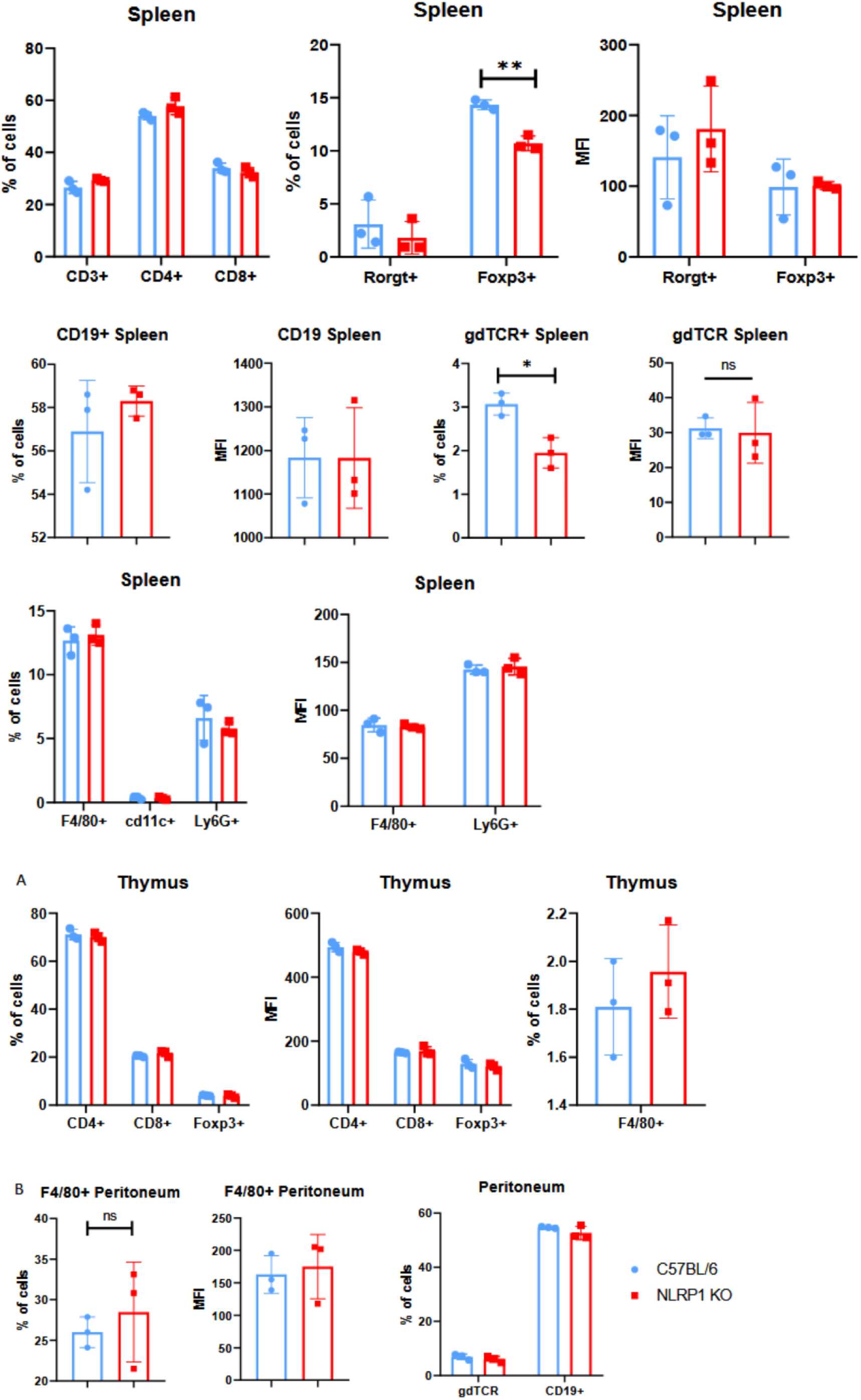
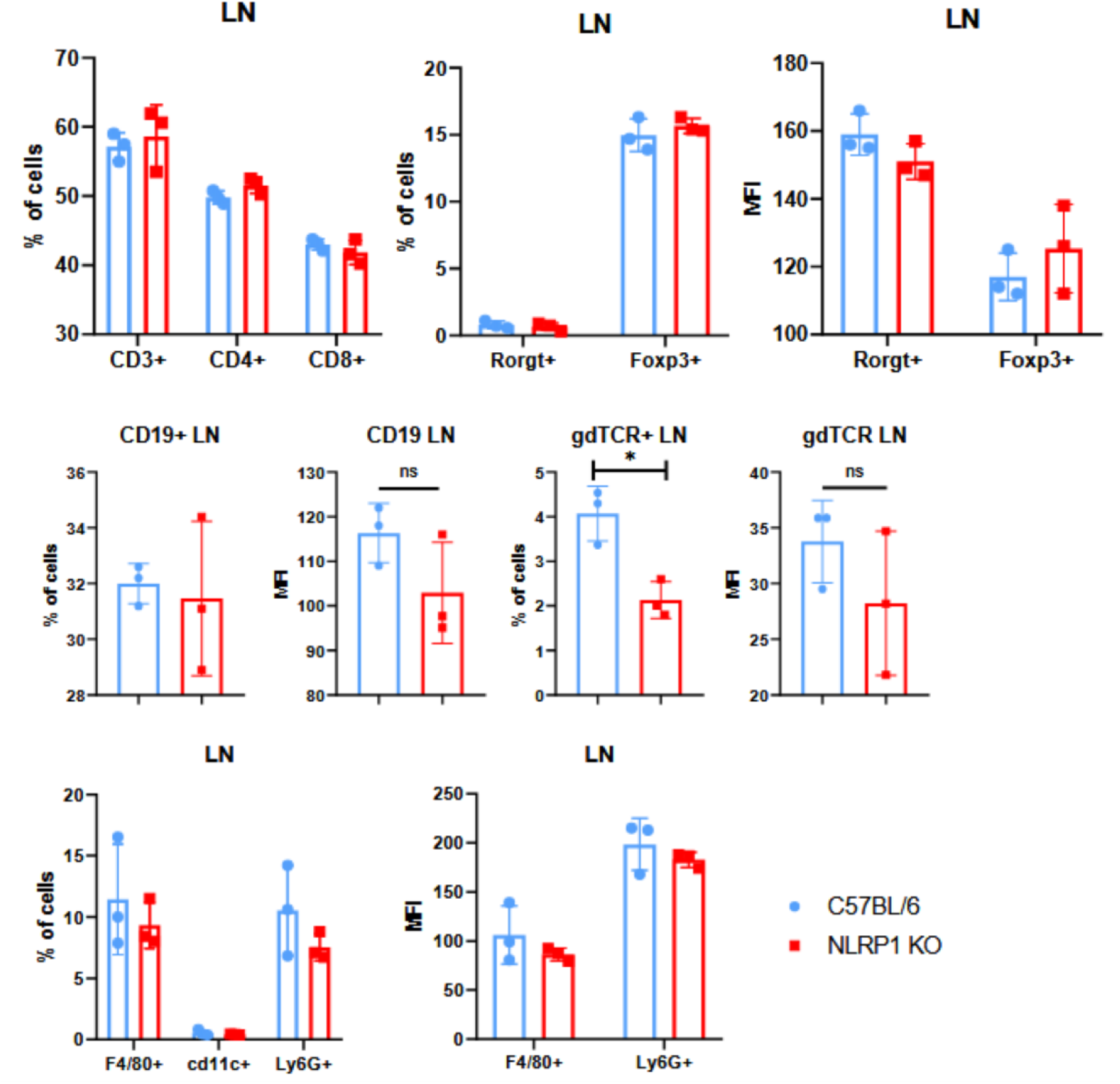
Immunophenotyping of C57BL/6 and NLRP1 KO mice was performed to assess the major immune cell populations in peripheral lymph nodes, spleen, thymus, and peritoneum. Lymphoid organs were isolated and subsequently analyzed for the main cell subtypes related to T cells, B cells, and γδ T cells, as well as innate subsets, based on surface molecule staining and the transcription factors RORγt and Foxp3, which are associated with Th17 and Treg subtypes, respectively. Foxp3 and RORγt expression were evaluated within the CD4^+^ T cell population. Data are presented as percentages of total cells and as median fluorescence intensity (MFI). Legend: * p < 0.05; ** p < 0.01;

**Sup. Figure 2.**
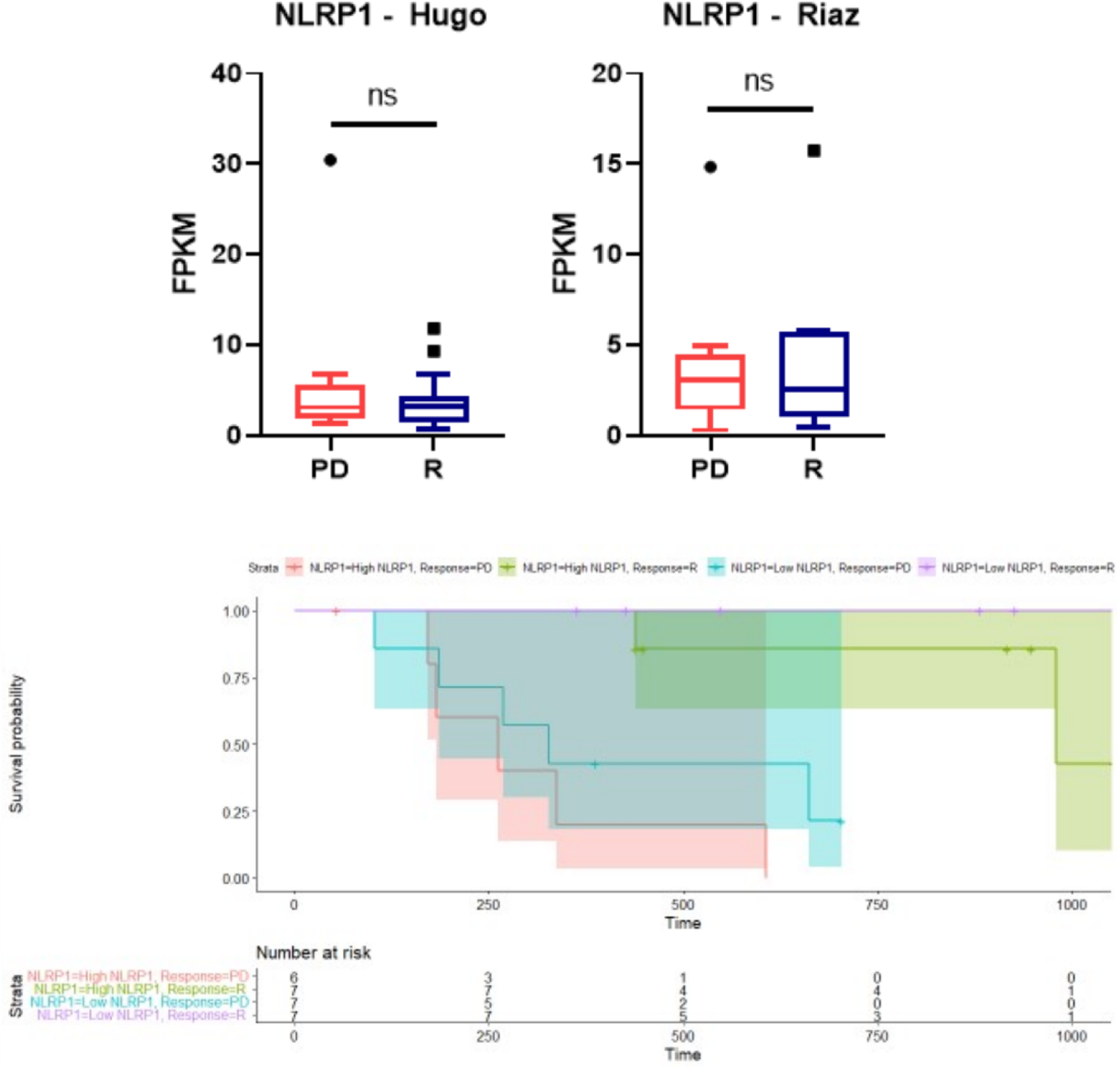
Relative expression of NLRP1 in FPKM among patients with progressive disease (PD) and responders (R) to immunotherapy, and its correlation with survival. (Upper) In both datasets, patients were classified as responders or non-responders and analyzed for NLRP1 expression levels. (Lower) Survival curve showing the relationship between anti–PD-1 response and NLRP1 expression levels, dichotomized by the median expression value in the Hugo dataset. A multivariable Cox regression analysis was performed. Legend: ns, not significant.

**Sup. Figure 3.**
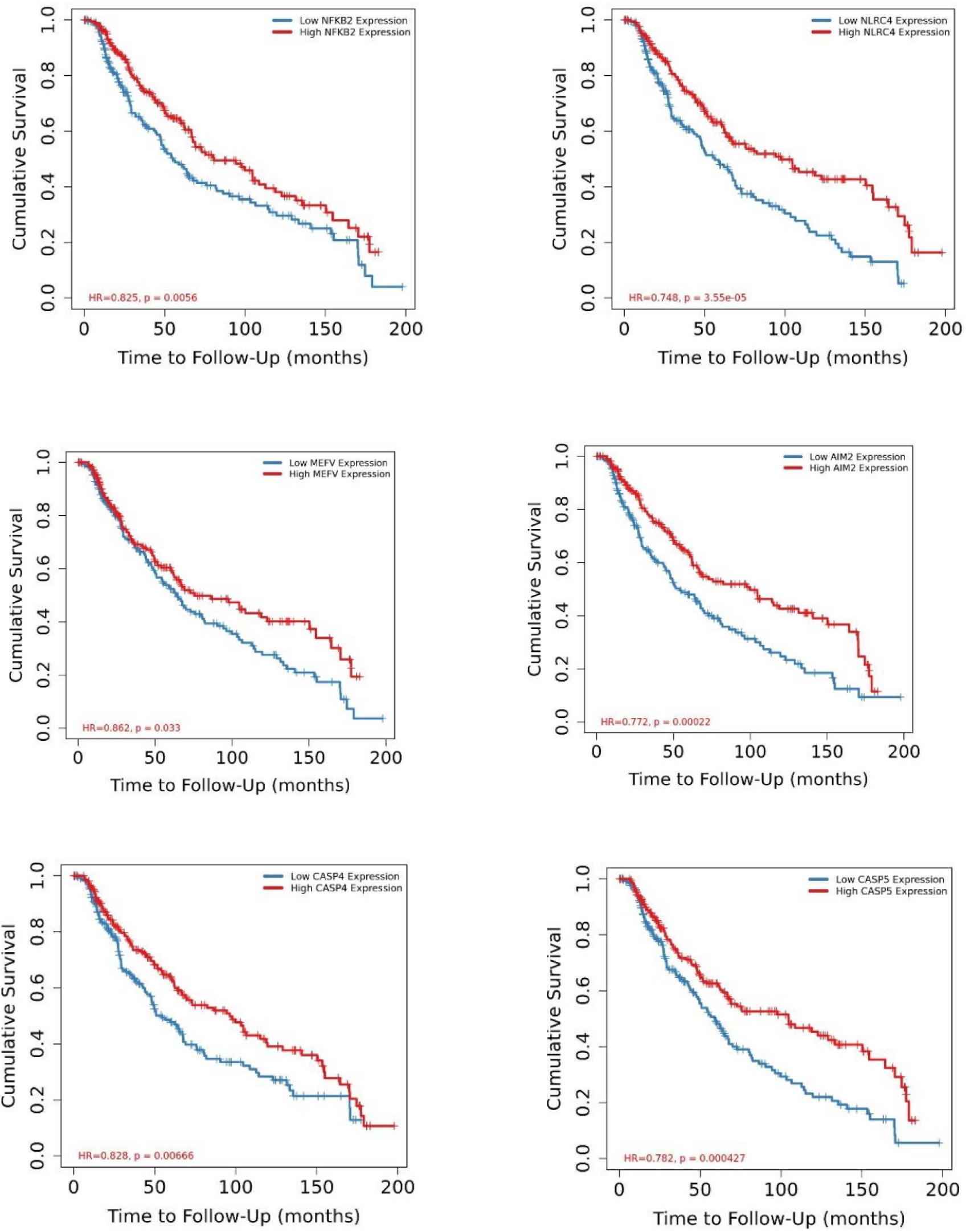
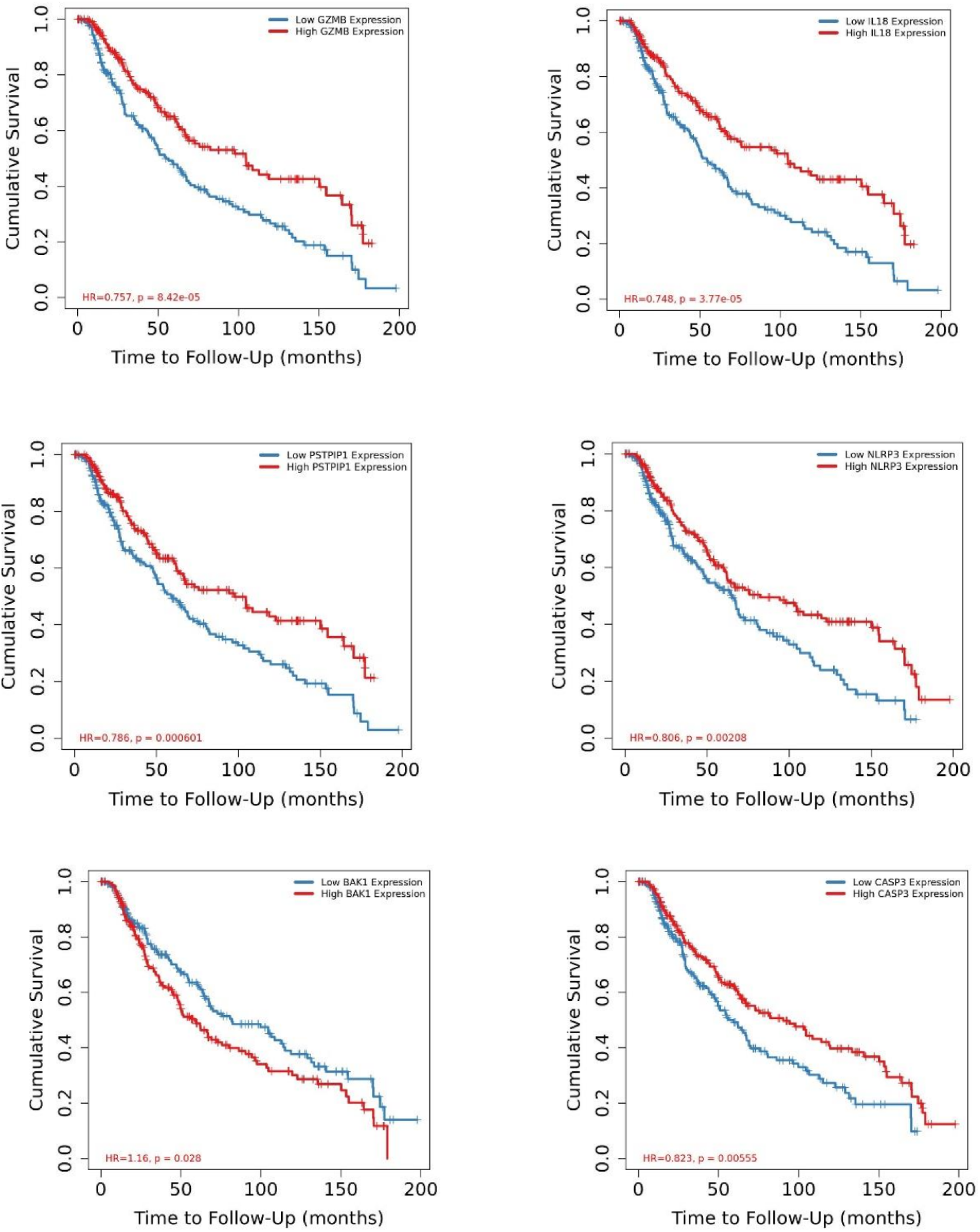
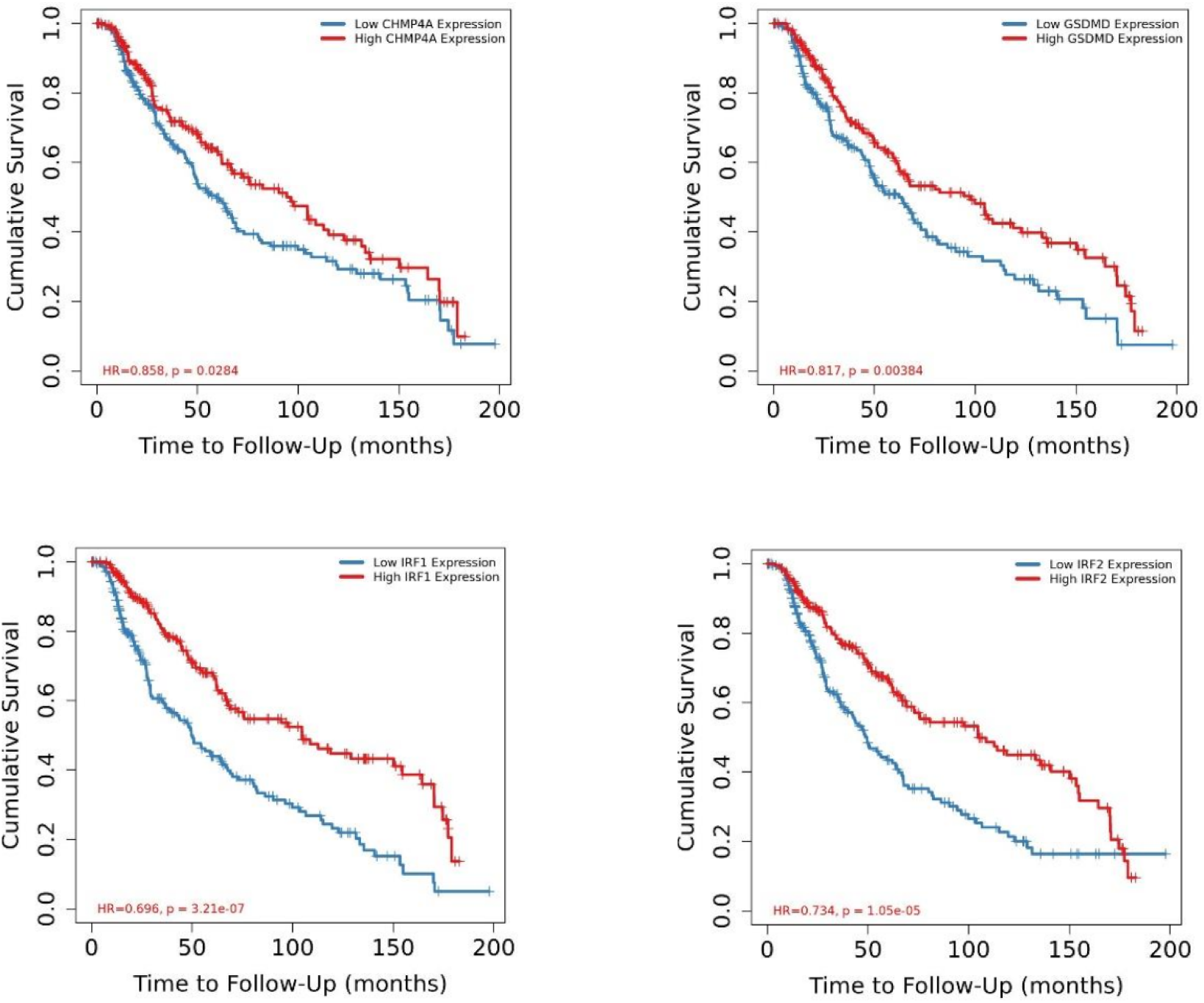
Survival curves of patients from the TCGA-SKCM cohort based on the genes used to construct the risk score.

**Sup. Figure 4.**
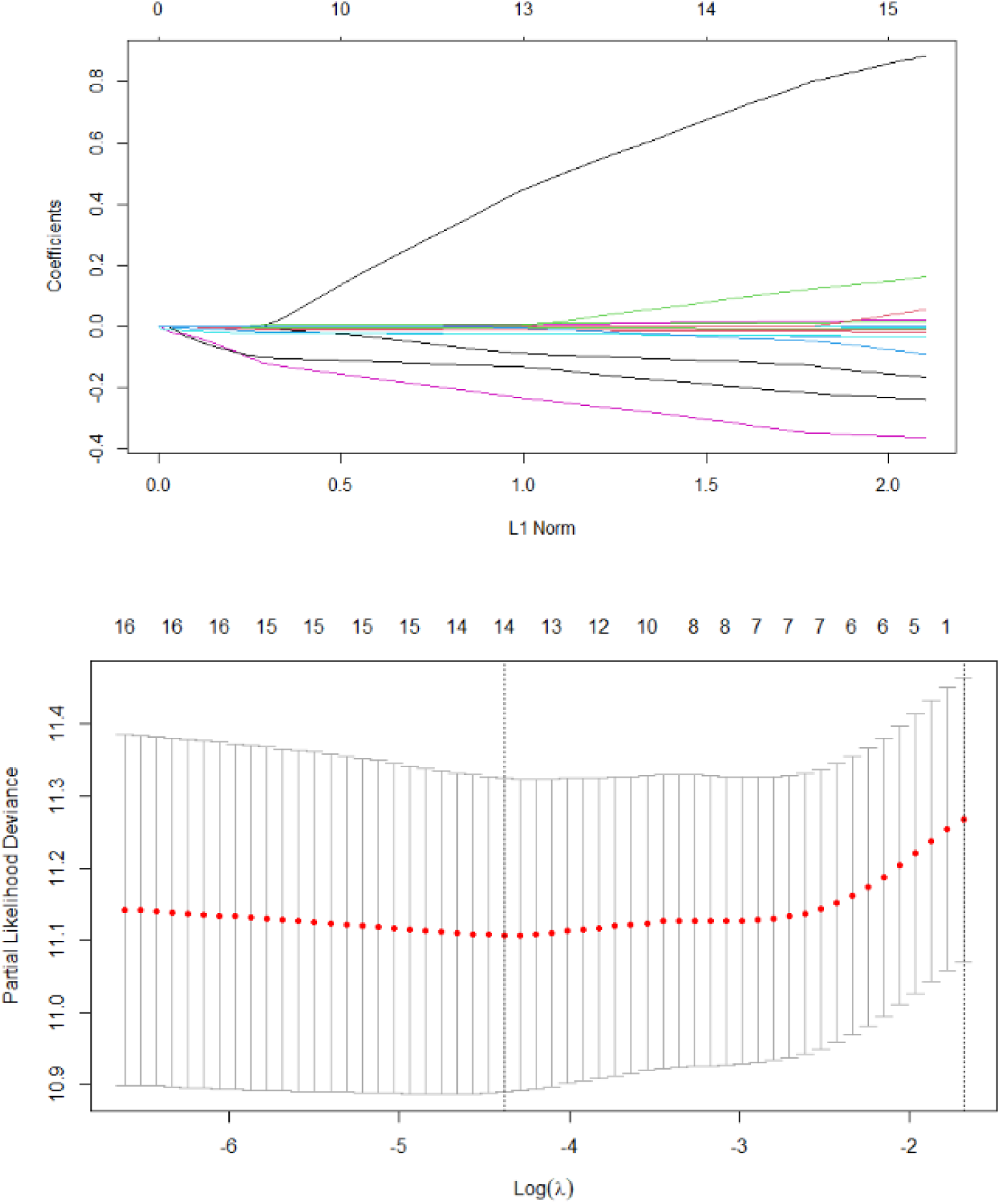
Construction of a risk score based on inflammasome- and pyroptosis-related genes. (Upper) Plot of the LASSO-Cox model fitting process, showing the LASSO coefficient for each gene that was effectively associated with patient survival. (Lower) Determination of the minimum lambda value in relation to the number of genes selected by the LASSO regression through cross-validation. Fourteen genes were retained at the lambda that minimized the cross-validation error.

